# Comparative genomics and community curation further improve gene annotations in the nematode *Pristionchus pacificus*

**DOI:** 10.1101/2020.08.03.233726

**Authors:** Marina Athanasouli, Hanh Witte, Christian Weiler, Tobias Loschko, Gabi Eberhardt, Ralf J. Sommer, Christian Rödelsperger

## Abstract

**Background:** Nematode model organisms such as *Caenorhabditis elegans* and *Pristionchus pacificus* are powerful systems for studying the evolution of gene function at a mechanistic level. However, the identification of *P. pacificus* orthologs of candidate genes known from *C. elegans* is complicated by the discrepancy in the quality of gene annotations, a common problem in nematode and invertebrate genomics.

**Results:** Here, we combine comparative genomic screens for suspicious gene models with community-based curation to further improve the quality of gene annotations in *P. pacificus*. We extend previous curations of one-to-one orthologs to larger gene families and also orphan genes. Cross-species comparisons of protein lengths and screens for atypical domain combinations and species-specific orphan genes resulted in 4,221 candidate genes that were subject to community-based curation. Corrections for 2,851 gene models were implemented in a new version of the *P. pacificus* gene annotations. The new set of gene annotations contains 28,896 genes and has a single copy ortholog completeness level of 97.6%.

**Conclusions:** Our work demonstrates the effectiveness of comparative genomic screens to identify suspicious gene models and the scalability of community-based approaches to improve the quality of thousands of gene models. Similar community-based approaches can help to improve the quality of gene annotations in other invertebrate species, including parasitic nematodes.

## Background

The nematode *Pristionchus pacificus* was initially introduced as a satellite model organism for comparing developmental processes to *Caenorhabditis elegans* [1, 2]. More recently, it has emerged as an independent model organism for studying the genetics of phenotypic plasticity [3–5] and behavior [6–8], interactions between host and microbes [9–11], and genome evolution [12–14]. Central to all these studies was the genome sequence of *P. pacificus*, which has undergone continuous improvements over time [15–17]. However, until recently, its gene annotations were almost exclusively based on automated pipelines that combined gene predictions and evidence-based annotations [18–20]. As a consequence, the gene annotations of *P. pacificus* did not match the quality of the highly curated *C. elegans* genome. This made it difficult for researchers from the *C. elegans* field to adapt *P. pacificus* for comparative studies, even though the availability of genetic toolkits including transgenic reporter lines and gene knockouts makes *P. pacificus* ideally suited for comparative studies of gene function [8, 21, 22]. Therefore, we have recently started to combine comparative genomic screens for suspicious gene models with community-based manual curation to improve the quality of the gene annotations in *P. pacificus* [23]. This pilot study screened for missing one-to-one orthologs of *C. elegans* genes in *P. pacificus*. Community-based curation of these candidate gene loci resulted in a substantial improvement of the *P. pacificus* gene annotations (version: El Paco annotation V2). Precisely, when assessed by benchmarking of universally conserved single copy orthologs (BUSCO) [24], the completeness level increased from 86% to 97%. Most missing orthologs were due to fused gene models some of which had long untranslated regions (UTRs) that actually contained complete genes. These errors could be corrected by manual inspection of the suspicious gene loci under the consideration of two recent transcriptome assemblies that were generated from strand-specific RNA-seq [25, 26] and Iso-seq data [27].

Here, we employ comparative genomic approaches to screen for further errors in other gene classes including large gene families that have undergone lineage-specific duplications [28] and species-specific orphan genes (SSOGs) [29] that were not the focus of our previous study [23]. Candidate loci are then curated by community-based manual inspection and eventually, corrections were proposed mainly based on available transcriptome assemblies. Overall, we investigated 4,221 suspicious gene models and implemented 2,851 corrections. This resulted in a further improved set of gene annotations for *P. pacificus*. Similar community-based curation approaches can help approving gene annotations in other nematode genomes including those of animal and plant parasites [23].

## Results

### Protein length comparison of orthologs identify hundreds of suspicious gene models

In our previous study, we focused on the identification of missing one-to-one orthologous genes in the *P. pacificus* genome and the identification of artificial fusions between two adjacent *P. pacificus* genes both of which have one-to-one orthologous genes [23]. Here, we aim to further improve the quality of one-to-one orthologous genes by finding and curating *P. pacificus* genes that are either unusually large and or small with regard to their *C. elegans* counterpart. We performed a comparison of protein length of 8,348 one-to-one orthologs between *C. elegans* and *P. pacificus* (Fig. 1a-c). Protein lengths between one-to-one orthologs are well correlated (Pearson’s r=0.83, Fig. 1a). However, there are slight differences in the length distributions (Fig. 1b,c) and using an arbitrary cutoff of a two-fold difference in protein length, we defined 532 *P. pacificus* genes as candidates for manual inspection. For example, in the case of the *P. pacificus* gene PPA00494 (ortholog of *C. elegans lev-8*), its predicted protein sequence encompasses 1094 amino acids, which is more than twice as long as *C. elegans* LEV-8 (531 amino acids) (Fig. 1d). Also, BLASTP analysis against the *C. elegans* proteins (version WS277) shows that the N-terminal part of PPA00494 is homologous to another *C. elegans* protein, Y73B6BL.37 (Fig. 1d), suggesting that it could represent an artificial gene fusion. Subsequent inspection in the genome browser showed two transcripts that were assembled from strand-specific RNA-seq data [26], which span the PPA00494 locus (Fig. 1e). This strongly supports that PPA00494 should be split by replacing it with the two assembled transcripts. After community-based curation, 309 (57%) corrections were proposed. The remaining cases were judged as either inconclusive (due to the lack of transcriptomic support) or correct. These results demonstrate that protein length comparisons between one-to-one orthologs are an effective way to identify suspicious gene models and to further improve the quality of one-to-one orthologs.

**Fig. 1.**
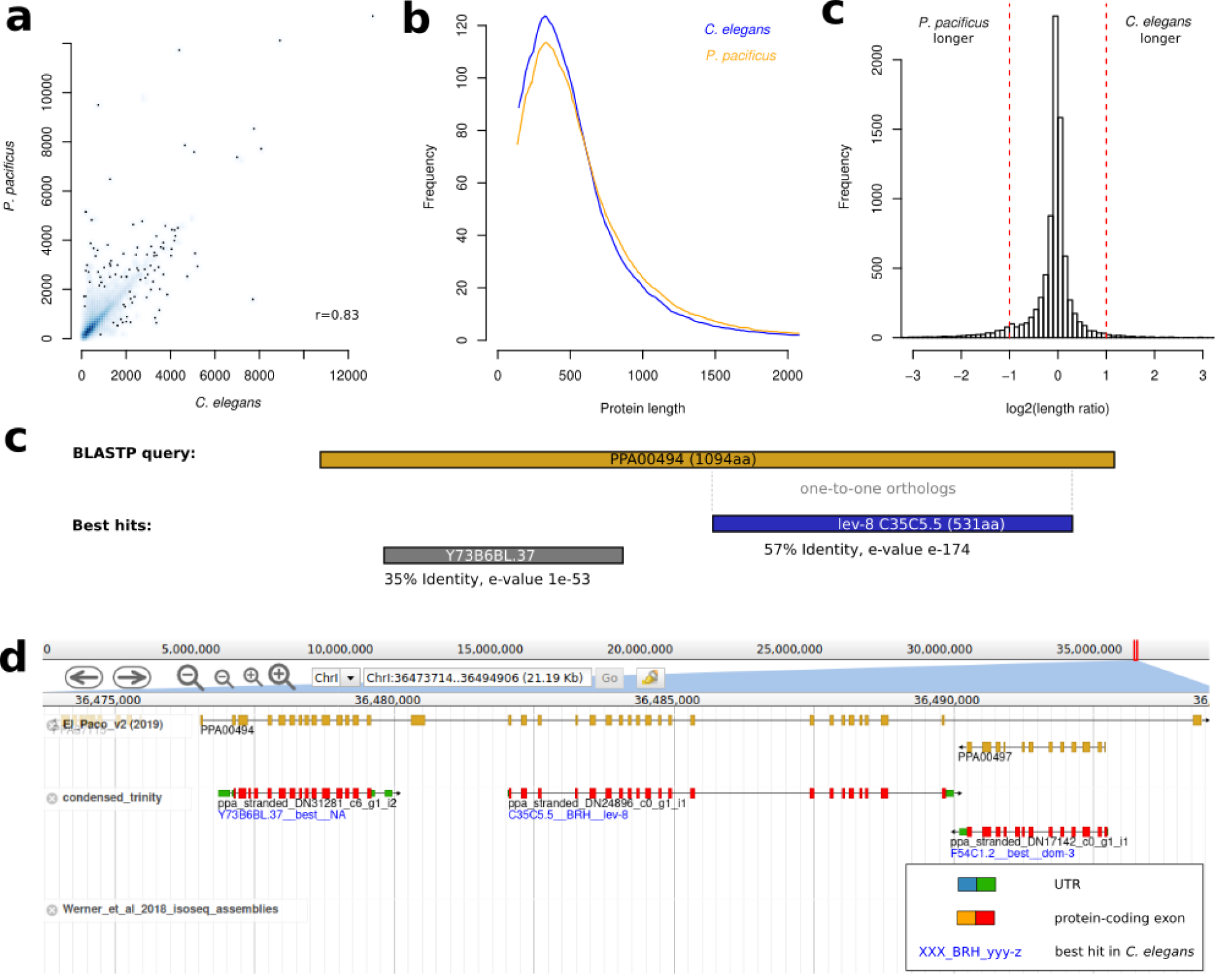
Comparison of protein lengths between one-to-one orthologs. **a** One-to-one orthologous genes between *C. elegans* and *P. pacificus* have highly similar protein lengths (Pearson’s r=0.83). **b** Size distributions of one-to-one orthologs show a peak at around 300 amino acids.**c** *P. pacificus* genes with more than two-fold length difference were considered for manual curation. **c** The *P. pacificus* one-to-one ortholog (PPA0494) of *C. elegans lev-8*, is more than twice as long as LEV-8. BLAST analysis showed that the N-terminal region has similarity to another *C. elegans* gene (Y37B6BL.37) suggesting that it represents an artificial gene fusion. **d** Manual inspection of the PPA0494 in the genome browser shows that there are two assembled RNA-seq transcripts (red) that cover most of the original gene model and further support that PPA0494 is an artificially fused gene model.

### Analysis of protein domains identifies further artificial gene fusions

Our previous study showed that the combination of incorrectly predicted gene boundaries and overlapping UTRs between neighboring genes in regions with high gene density most likely caused artificial gene fusions. In order to screen for further cases of artificial gene fusions, we applied a comparative approach to identify proteins with atypical domain combinations that do not exist in other nematodes such as *C. elegans*, and more distantly related *Bursaphelenchelus xylophilus* [30], and *Strongyloides ratti* [31]. This yielded 1,589 *P. pacificus* candidates (Table 1) for further inspection. Note, that such atypical domain combinations are not necessarily artifacts. For example, the same screen in the highly curated *C. elegans* genome, identified 932 genes with atypical domain combinations. Manual inspection of these gene models and available transcriptome assemblies in the WormBase genome browser (WS177) combined with BLASTP analysis against *C. briggsae* revealed three candidates for putatively incorrect annotation in *C. elegans*, which deserve closer inspection (Additional file 1, Figure S1). After community curation of the *P. pacificus* candidates, corrections were proposed for 695 (44%) candidates. Next, we defined 1,325 unusually small or long members of 25 highly abundant gene families as further candidates for manual inspection (Fig. 2a). After community curation, corrections were proposed for 420 (32%) of these candidates. The three described screens partially identify the same candidates (Fig. 2b), yet the presence of hundreds of candidate genes that are specific to each method indicates how complementary these different approaches are.

**Table 1.** Comparative assessment of different *P. pacificus* gene annotations

| Category | *P. pacificus* El Paco gene annotations |  |
| --- | --- | --- |
|  | V2 | V3 |
| Number of genes | 28,036 | 28,896 |
| Protein-coding sequence (Mb) | 35.3 | 35.3 |
| BUSCO Completeness (%) | 97.1 | 97.6 |
| BUSCO Duplicated (%) | 1.7 | 1.8 |
| BUSCO Fragmented (%) | 2.0 | 2.0 |
| BUSCO Missing (%) | 0.9 | 0.4 |
| Number of 1-1 orthologs (BRHs) | 8,348 | 8,607 |
| Number of 1-1 orthologs with variable protein length (%) | 532 | 265 |
| Number of proteins with atypical domain combinations | 1,589 | 1,137 |
| Number of protein family length outlier | 1,325 | 1,201 |

**Fig. 2.**
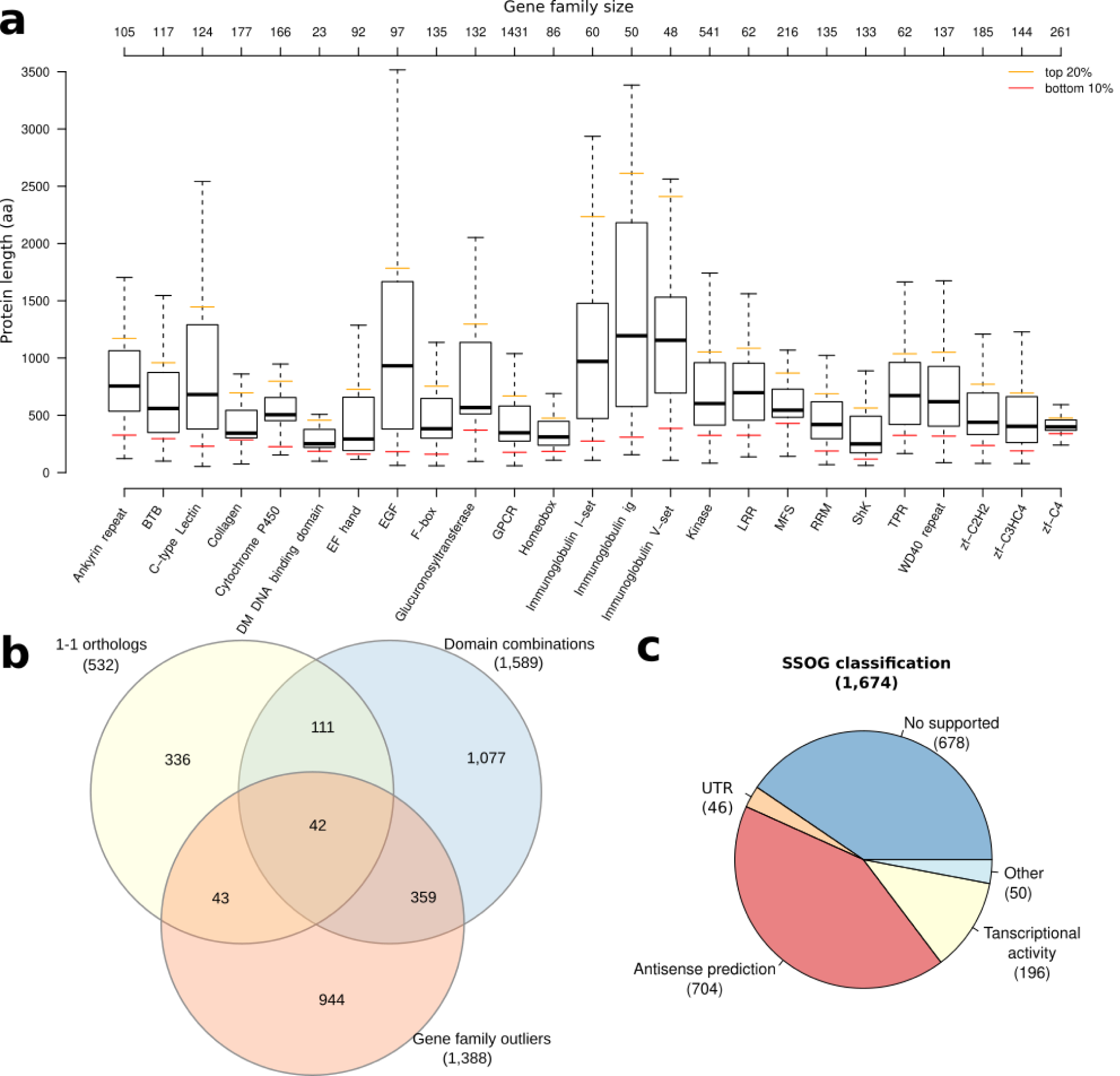
Identification of candidates for manual curation. **a** The boxplots show the length distributions of members of 25 highly abundant gene families. The lower 10% and the upper 20% of each gene family were selected for manual inspection. **b** Individual screens for suspicious gene models reveal between 336 to 1077 specific candidates indicating their highly complementary. **c** Manual classification of *P. pacificus* SSOGs shows numerous genes that overlap gene models on the opposite strand.

### Gene prediction artifacts are a likely source of SSOGs

A previous analysis of *P. pacificus* orphan genes revealed that the majority of SSOGs had no transcriptomic support [14]. Based on reanalysis of the current gene annotations with available phylogenomic and phylotranscriptomic data [26, 32], we identified 1,988 (7%) *P. pacificus* SSOGs of which 314 were classified as having transcriptomic support. Manual inspection of the remaining SSOGs classified 678 (41%) of candidates as not having any transcriptional support (Fig. 2c), even when considering additional transcriptomic data sets such as iso-seq or dauer-specific transcriptomes [27, 33]. Further 196 (12%) of SSOG candidates showed some transcriptional activity, but this expression data was mostly not sufficient to support their gene structure. Strikingly, we found 704 (42%) and 46 (3%) SSOGs, which overlapped existing gene models on the antisense strand of protein-coding exons and UTRs, respectively (Fig. 2c and 3a,b). However, visual inspection of transcriptomic data only supported the sense gene as opposed to the antisense SSOG. As there is neither protein homology nor transcriptional data supporting these antisense SSOGs, we would tend to argue that these spurious antisense gene models most likely derive from the contribution of gene prediction softwares SNAP and AUGUSTUS during the process of the original gene annotation [17, 19, 20]. In total, we removed 1,515 of these unsupported SSOGs, as their lack of transcriptional evidence makes it difficult to conclusively study the process of novel gene formation [14, 29].

**Fig. 3.**
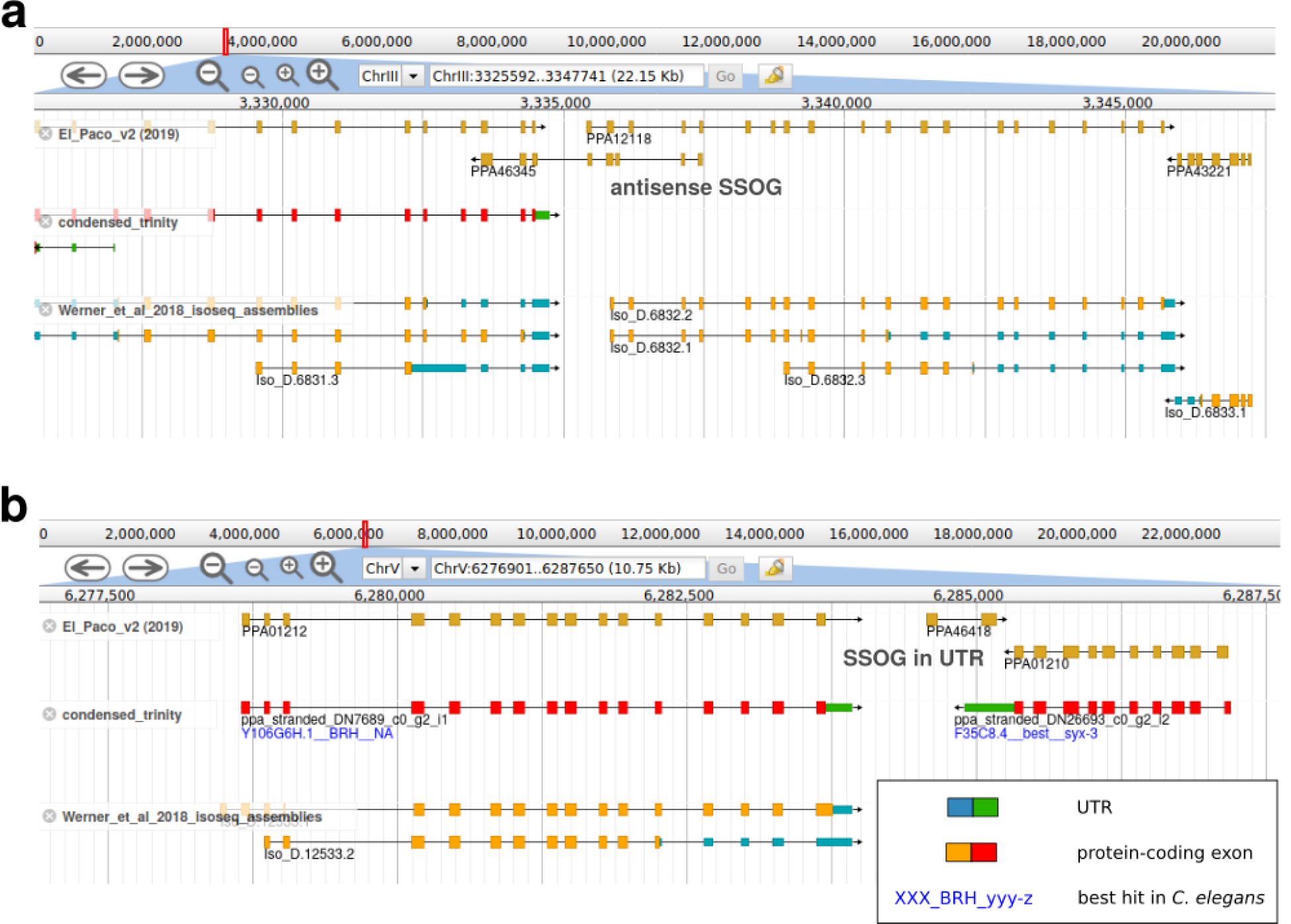
Examples of unsupported SSOGs. **a** The *P. pacificus* SSOG PPA46345 overlaps exons of two other gene models that are well supported by transcriptome assemblies from strand-specific RNA-seq and Iso-seq data. **b** The *P. pacificus* SSOG PPA4618 overlaps the UTR of a well supported gene model. The absence of strand-specific transcriptomic support indicates that *P. pacificus* SSOGs PPA46345 and PPA4618 are likely gene prediction artifacts.

### New *P. pacificus* gene annotations show increased homogeneity and better reflect existing RNA-seq data

In total, we visually inspected 4,221 suspicious gene models and proposed corrections for 2,851 (68%). We implemented all proposed corrections into a new *P. pacificus* gene annotation (version: El Paco gene annotation V3), which comprises 28,896 gene models and spans 35.2 Mb of protein-coding sequence with a BUSCO completeness level of 97.6% (Table 1). As expected, the numbers of one-to-one orthologs with length differences, the number of genes with atypical domain combinations, and the number of gene family outliers went down by 10-50%. To additionally test if the new set of gene annotations better captures RNA-seq data sets, we reanalyzed 15 RNA-seq data sets from four different studies [9, 13, 34, 35] and quantified the percentage of reads that could be assigned to features of the gene annotations. The new set of gene annotations consistently captures two percent more of the RNA-seq alignments (Table 2). Considering that the total amount of annotated protein-coding sequence remained almost unaltered, this supports that the new set of gene annotations better reflects RNA-seq data.

**Table 2.** Percentage of assigned RNA-seq reads to different sets of gene annotations.

| *P. pacificus* RNA-seq samples |  | Successfully assigned alignments (%) |  | Reference |
| --- | --- | --- | --- | --- |
| Accession | Description | V2 | V3 |  |
| ERR777792 | Mixed-stage on *E. coli* OP50 | 74.8 | 76.8 | [13] |
| ERR777793 | Mixed-stage on *E. coli* OP50 | 74.9 | 76.6 | [13] |
| ERR777794 | Mixed-stage on *E. coli* OP50 | 74.4 | 76.1 | [13] |
| SRR4017216 | Adults on *E. coli* OP50 | 79.8 | 81.7 | [34] |
| SRR4017217 | Adults on *E. coli* OP50 | 80.3 | 82.2 | [34] |
| SRR4017218 | Adults on *Cryptococcus* C3 | 79.6 | 81.6 | [34] |
| SRR4017219 | Adults on *Cryptococcus* C3 | 79.2 | 81.1 | [34] |
| SRR4017220 | Adults on *Cryptococcus* C5 | 79.9 | 81.8 | [34] |
| SRR4017221 | Adults on *Cryptococcus* C5 | 80.7 | 82.6 | [34] |
| ERR3421261 | Adults on *E. coli* OP50 | 79.7 | 81.6 | [9] |
| ERR3421262 | Adults on *E. coli* OP50 | 79.5 | 81.3 | [9] |
| ERR3421263 | Adults on *Novosphingobium* L76 | 79.6 | 81.5 | [9] |
| ERR3421264 | Adults on *Novosphingobium* L76 | 79.5 | 81.5 | [9] |
| SRR2142256 | Adults on *E. coli* OP50 | 77.8 | 79.8 | [35] |
| SRR2142257 | Intestines | 72.5 | 74.2 | [35] |

## Discussion

In the early genomic era, gene annotation was heavily dependent on automated gene finding algorithms that tried to recognize gene structures based on statistical sequence properties of exons, introns, and splicing sites [19, 20]. This was highly suited when functional data, e.g. expressed sequence tags and cDNAs, were scarce and the only way to annotate a complete genome was to extract informative sequence features from a limited test set and extrapolate them to the whole genome. However, with the dramatic improvement of sequencing protocols and technologies, it became feasible to generate evidence-based gene annotations from transcriptome and homology data [18, 36]. Under the consideration that related genomes at an optimal evolutionary distance to a focal organism and transcriptomic evidence for all genes are rarely available, this still justifies the usage of gene prediction tools. In the case of the *P. pacificus*, previous versions of gene annotations that were completely based on the results of gene prediction tools were suited to perform evolutionary genomic analysis and genetic screens [37, 38]. Subsequently, we employed the widely used MAKER2 pipeline to generate a more comprehensive gene annotation by integration of large-scale transcriptomic and protein homology data as well as gene predictions [17, 36]. Comparative analysis of genome quality for 22 nematode species revealed that already these gene annotations (version: El Paco annotation V1) were of relatively high quality (86% BUSCO completeness) [23]. Nevertheless, the question of how good gene annotations need to be will depend on what researchers want to do with them. Reverse genetic studies in nematodes with well established genetic toolkits are extremely powerful systems for comparative studies of gene function [8, 22] and the evolution of the nervous system and associated behaviors [39, 40]. Yet, the identification of *P. pacificus* orthologs for candidate *C. elegans* genes with known function is complicated by the widespread abundance of lineage-specific duplications [28, 33], but also by the difference in the quality of gene annotations. Facilitating the easy adaptation of *P. pacificus* as a comparative model system for *C. elegans* researchers, who are used to working with one of the best and well-characterized genomes, is one of our main motivations for this study. The chromosome-scale assembly of *P. pacificus* has already been a major step to minimize the disparity between the genomic resources of both species [17]. Lifting up the quality of gene annotations to a comparable level will thus further increase the attractiveness of the *P. pacificus* system for evolutionary studies.

Another motivation for continuous efforts in improving the quality of gene annotations is our focus on the origin and evolution of orphan genes in *P. pacificus* [41–43]. Initially, around one-third of the *P. pacificus* gene repertoire was defined as orphan genes without homology in the genomes of other nematode families [16, 37]. Unbiased genetic screens have identified orphan genes that control important biological processes such as developmental decisions and predatory behavior [6, 42]. Phylogenomic investigation of ten diplogastrid genomes revealed the evolutionary dynamics of these novel genes and built the framework to dissect the diversity of mechanisms of origin [14, 32]. When we screened for high quality SSOG candidates for origin analysis, we found that the majority of SSOGs had no transcriptomic support. Together with the finding that SSOGs constitute an unusually large age class (phylostratum), this made us wonder to what extent this gene class might possibly be inflated by gene annotation artifacts [14]. Therefore, we revisited 1,674 candidates and confirmed that most of them indeed show no evidence of transcription. In addition, we found 704 SSOGs, which overlapped other gene models on the antisense strand and whenever available, strand-specific RNA-seq did not support the SSOG gene model. Even though it is expected that SSOGs show little evidence of expression and we cannot conclusively argue that these gene models are gene annotations artifacts (they represent coding potential that might be used under some conditions), for practical reasons we chose to remove them to allow future investigations of orphan origin to start with a set of well supported candidate SSOGs. Thus, we hope that the community-based curation of the *P. pacificus* gene annotations will help future studies in many aspects of evolutionary biology.

## Conclusions

Our work demonstrates that even for non-classical model organisms with small research communities, manual inspection and curation of thousands of genes can be achieved. Thereby numerous comparative genomic screens can be applied to enrich the candidate set for suspicious gene models that actually need to be corrected. The example of the highly curated *P. pacificus* genome emphasizes the effectiveness and scalability of manual curation for many other genome projects including those of nematode animal and plant parasites.

## Methods

### Candidate identification based on length comparison of orthologous proteins

We obtained 8,348 one-to-one orthologs between *C. elegans* and *P. pacificus* that were predicted based on best reciprocal BLASTP hits in a previous study [23]. We then calculated the protein length ratio between the *P. pacificus* and *C. elegans* one-to-one orthologs. In case of multiple isoforms for a given gene, we chose the isoform with the longest protein sequence (WormBase release WS260). Based on an arbitrary cutoff of a two-fold difference in protein length between the two species, we identified 531 *P. pacificus* candidates for manual curation.

### Candidate identification based on protein domain content

We ran the hmmsearch program of the HMMER package (version 3.0, e-value < 0.001, profiles from PFAM-A.hmm) on protein sets of *C. elegans* (WS260), *P. pacificus* (El Paco V2), *B. xylophilus* (WS248), *S. ratti* (WS260). We counted occurrences of protein domains and defined as candidates, domain combinations that are unique to *P. pacificus* and occur at low frequencies (less than ten times). This yielded 1589 candidates with atypical protein domain combinations. Next, we selected 25 highly abundant gene families such as collagens and C-type lectins that were defined by a PFAM domain and classified further candidate proteins if their length fell under the first or above the eighth decile of the length distribution of all members of a given gene family. This identified 1,388 candidate genes for manual curation.

### Identification and curation of *P. pacificus* species-specific orphan genes

We defined *P. pacificus* SSOGs by BLASTP searches of the *P. pacificus* proteins (version: El Paco annotation V2) against annotated protein sets and predicted ORFs in assembled transcripts of *P. exspectatus, P. arcanus, P. maxplancki*, and *P. japonica* [26, 32]. This identified 1,988 (7%) *P. pacificus* SSOGs without a BLASTP hit in any of the reference data sets (e-value <0.001). 314 SSOGs showed transcriptomic support as they had a BLASTP hit in ORFs of the *P. pacificus* transcriptome assembly. The remaining 1,674 were defined as SSOGs without transcriptomic support and were thus considered as candidates for manual curation.

### Community-based manual curation of gene models

Community-based gene curation was performed as described in our pilot study [23]. In short, candidate lists were shared in online spreadsheets and individual genes were visually inspected in the jbrowse genome browser instance on http://www.pristionchus.org [44]. Based on available transcriptomic resources, which include RNA-seq data from different developmental stages, strand-specific transcriptome assemblies from mixed-stage cultures [25, 26], and iso-seq data [27], community curators were trained to evaluate whether a locus was well covered by transcriptomic data and in case of evidence for an artificial gene fusion to propose the replacement of the original gene model by assembled transcripts. If the genomic neighborhood of the candidate genes showed obvious inconsistencies between original gene models and transcriptome data, we eventually curated such neighboring genes. However, we omitted any gene that was curated in our previous study, as these changes were not yet fully implemented in the latest WormBase release WS177 of *P. pacificus* and we wanted to avoid version conflicts. While for most candidate genes, we did not propose any correction in case that available transcriptomic data was insufficient to make a conclusive statement, in the case of SSOGs, we removed the gene model if not at least some partial support RNA-seq data supported the gene structure (see *Discussion*).

### Quality assessment of gene annotations

In order to evaluate the quality of gene annotations, we ran the BUSCO program (version 3.0.1) in protein mode (option: -m prot) against the nematode_odb9 data set (N = 982 orthologs) [24]. To test whether the new set of gene annotations better captures RNA-seq data, we downloaded 15 RNA-seq data sets from the European Nucleotide Archive and aligned these data sets against the *P. pacificus* reference genome (version: El Paco) with the help of the STAR aligner (version: 2.5.4b, default options, reference was the *P. pacificus* genome without any gene annotation) [45]. Next, we quantified the percentage of alignments that could be assigned to gene annotations using the featureCounts function of the Rsubread library in R (version 4.0.0).

## Ethics approval and consent to participate

Not applicable

## Consent for publication

Not applicable

## Availability of data and materials

The new set of *P. pacificus* gene annotations (version: El Paco gene annotation V3) is publicly available at http://www.pristionchus.org/download/ and was also submitted to WormBase where it will be published following further curation.

## Authors’ contributions

Conceptualization, C.R.; Investigation, C.R., M.A., H.W., C.W., T.L. and G.E.; Data curation, C.R., M.A., H.W., C.W., T.L. and G.E.; Visualization, C. R.; Writing original draft, C.R.; Writing – review & editing, C.R., and R.J.S.; Project administration, C. R.; Supervision, C.R. and R.J.S.; Funding acquisition, R.J.S.

## Acknowledgements

The authors would like to thank the complete *Pristionchus* community for their long-term interest in studying *P. pacificus* and thus motivating this work.

