## Additional file 1, Figure S1 for "Comparative genomics and community curation further improve gene annotations in the nematode *Pristionchus pacificus*"

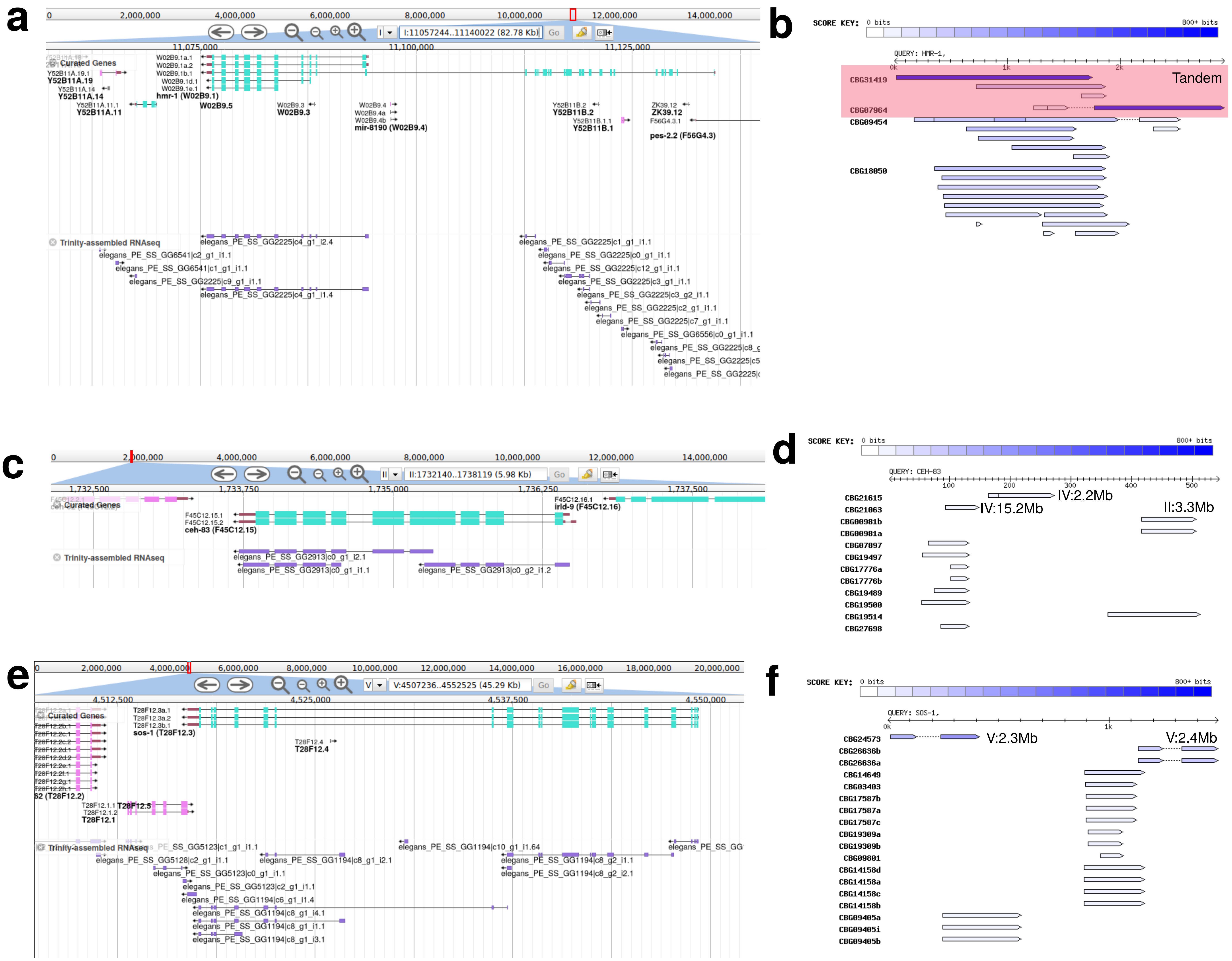


**Figure S1 Putative candidates of annotation errors in *C. elegans*.** 932 *C. elegans* candidates that were defined based on atypical protein domain combinations were manually inspected in the WormBase genome browser (WS177). **a** The longest isoform of the *C. elegans* gene *hmr-1* is not fully supported by transcripts that were assembled from RNA-seq data. **b** BLASTP analysis against the *C. briggsae* proteins identified two separate genes, located in a tandem configuration, that share partial matches with the longest isoform of *hmr-1*, suggesting potential annoation problems in either of the two species. **c** The gene *ceh-83* is also only partially supported by the transcriptome assembly. **d** BLASTP analysis identifies partial matches against multiple *C. briggsae* genes, which are scattered across the genome. Therefore, the gene model of *C. elegans ceh-83* could represesent an artificial gene fusion. **e,f** Incomplete support by transcriptome assemblies and BLASTP hits to separate *C. briggsae* proteins point towards potential annotation problems of *C. elegans sos-1*.
